# G-quadruplex sequence dictates tau oligomerization and fibril morphology

**DOI:** 10.64898/2025.12.02.691785

**Authors:** Lena M. Kallweit, Luke T. Kelly, Kevin M. Reynolds-Caicedo, Daniel Paredes, Scott Horowitz

## Abstract

Previous studies have identified that non-canonical nucleic acid structures known as G-quadruplexes (G4s) modulate protein aggregation and could play major roles in neurodegenerative diseases. Here we examine the presence and protein oligomerization activity of G4s that are enriched in human hippocampal aggregates. We found these G4s to be powerful, sequence-specific modulators of aggregation. Different G4s facilitated aggregate propagation in cells, caused tau to form distinct protein oligomer populations, and seeded tau fibrils with different fibril structure and length. This study highlights the importance of nucleic acid composition within aggregates, where small changes in nucleotide sequence and topology can vastly alter protein interactions, oligomerization, and fibrillization.

## 1. Introduction

Protein aggregation is a central pathological hallmark shared by many neurodegenerative diseases. [1–7] Initial observations in the late 1990s revealed that brain aggregates also harbor considerable levels of RNA and DNA, establishing nucleic acids as integral components of these aggregates. [8–10] These observations prompted testing whether nucleic acids could promote fibril formation of proteins involved in neurodegeneration, such as prion proteins and tau. Using nucleic acid sequences of unknown physiological relevance, it was shown that high concentrations of nucleic acids could play a role in the promotion of this aggregation *in vitro*. [11–14]

In 2021, Reis et al sequenced nucleic acids isolated from human brain aggregates. They demonstrated that these nucleic acids were enriched in polyguanine stretches with a high predicted propensity of forming G-quadruplex (G4) structures. [15] G4s are secondary structures formed in guanine-rich DNA and RNA sequences. Hoogsteen hydrogen bonding between four guanines stabilizes these stacked G-quartets. [16] G4s can form multiple structural topologies, including parallel, anti-parallel, and mixed 3+1 G4s, referring to the directionality of the nucleic acid strand (Fig 1A). Beyond overall topology, G4s contain a rich diversity of different detailed structures. [17] Both DNA and RNA G4s are often enriched in regulatory loci that affect replication, transcription, translation, and stability. [18–20] G4s are dynamic, folding and unfolding in response to their cellular environment. While they are actively unfolded under normal cellular conditions, they fold under stress and remain folded until the conditions have dissipated. [21]

**Figure 1.**
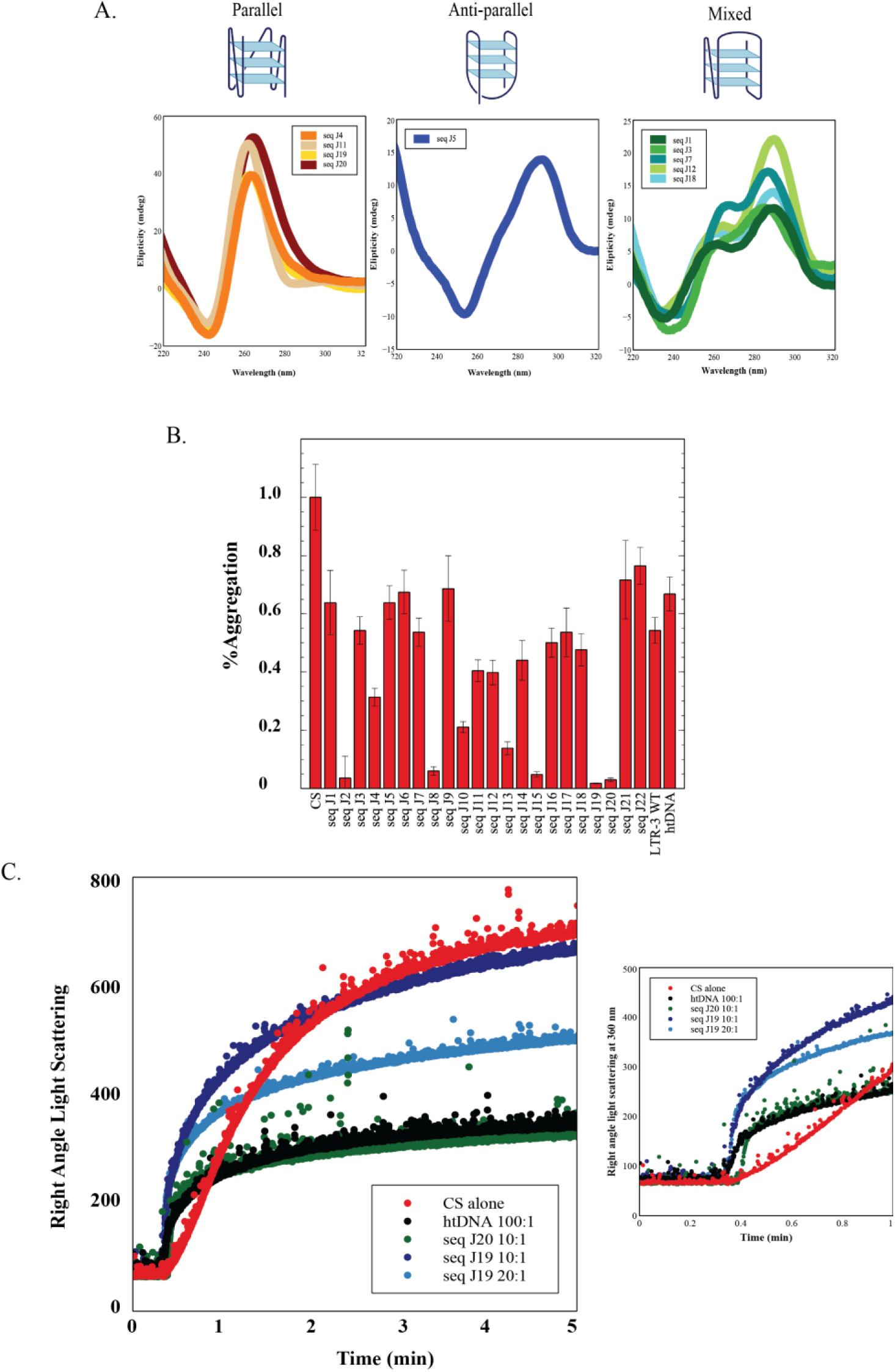
A) Circular dichroism of physiological sequences exhibits traces characteristic of parallel, anti-parallel, or mixed G-quadruplexes. B ) Aggregation of citrate synthase under heat stress when incubated with nucleic acid sequences compared to protein alone. C) Aggregation of denatured citrate synthase via light scattering when incubated with two different sequences or bulk DNA at differing concentrations as compared to protein alone. Zoom in reveals a steeper kinetic slope, indicating smaller oligomer formation.

Reis et al’s observation of G4s being enriched in human brain aggregates was particularly interesting because of concurrent work by several labs showing that G4s are extremely powerful modulators of protein folding and aggregation. G4s can catalyze protein folding [22] and aid the protein folding environment of the cell, [23, 24] as well as prevent protein aggregation *in vitro*. [25–27] On the other hand, G4s can also promote the formation of small protein oligomers [25, 26] and scaffold protein aggregation of disease-relevant proteins. For example, G4s interact with and in some cases promote the aggregation of proteins such as FMRP, α-synuclein, TDP-43, Huntingtin and tau. [11, 15, 28–34] G4s are enriched in the hippocampus in older individuals and those with more severe Alzheimer’s disease, where they colocalize with phospho-tau tangles. [35] G4 levels in the human brain are highly correlated to APOE allele type, with individuals with a high risk of Alzheimer’s Disease carrying higher levels of G4s in their hippocampus. [35]

These observations suggest G4 involvement in protein aggregation and neurodegenerative disease. However, the studies thus far on G4s in protein aggregation and related neurodegeneration have mainly used model G4 sequences of unknown physiological relevance. Effects vary based on G4 sequence and structure, suggesting that studying the physiologically relevant G4s individually could be particularly important. [25, 36] We therefore examined the mechanism of action of potential G4s enriched in human brain aggregates in modulating protein aggregation generally, and with a disease relevant protein, tau.

## 2. Methods

### Circular Dichroism

CD spectra were obtained using a Jasco J-1100 circular dichroism at 25°C. Sequence was resuspended in 10mM potassium phosphate pH 7.5 buffer and diluted to 25 µM (per strand) RNA. The CD measurement was taken from 320 nm to 220 nm at 1 nm intervals using a 50 nm/min scanning speed. Secondary structure was qualitatively interpreted as in [37].

### N-methyl-mesoporphyrin IX fluorescence

Fluorescence was measured in a clear-bottom 96-well plate (Corning). Each well contains 1 μM NMM with 5 μM annealed nucleic acid in 10 mM potassium phosphate buffer at pH 7.5. Emission spectra were measured at 24 °C using an excitation wavelength of 400 nm and an emission wavelength of 610 nm on a Tecan M200. Fluorescence is displayed as a ratio compared to NMM alone.

### Non-denaturing gel of RNA sequences

All oligonucleotides were ordered from IDT and annealed on the thermomixer before the following assays. The anneal program consists of 2 min at 95° C then cooling to 25° C at a ramp-down rate of 1° C/min. An estimated 500 ng/µL nucleic acid was loaded into a 20% TBE gel. Gels were stained with NMM, then SYBR gold.

### Thermal aggregation of citrate synthase

22 DNA sequences were incubated with Citrate synthase from porcine heart (Sigma-Aldrich C3260-5KU) at a 2:1 DNA to protein ratio in a Corning 3880 plate in 10mM Potassium Phosphate buffer, pH 7.5, as previously described [25]. Plates were transferred from ice to a preheated 50° C plate reader and run for 1.5 hours. Percent aggregation was calculated relative to maximum protein alone absorbance value. Error bars represent standard error propagated from triplicate protein alone and triplicate experimental measurement. Herring testes DNA (Sigma) was run as a control to ensure consistency across experiments.

### Chemical aggregation light scattering

RALS assay was performed as previously described. [25, 38] 12 µM Citrate Synthase was denatured in 6 M guanidine-HCl, 40 mM HEPES, overnight at 25°C. The denatured Citrate Synthase was diluted to 75 nM into 40 mM HEPES, pH 7.5 (KOH) the following morning in the presence of nucleic acids at different concentrations. Aggregation was measured using right-angle light scattering at 360 nm in a cuvette with consistent mixing.

### Expression and purification of tau constructs

All constructs were cloned into pet28b vectors with T7 promoters. 0N4R vector contains a HIS tag, T7 tag and thrombin site. Expression and purification protocols adapted from previous protocols. [13, 39] All constructs used were expressed in BL21(DE3)pLysS competent cells with 50 μg/mL working concentration of Kanamycin used throughout. Single colonies were grown in overnight small LB cultures, then transferred to larger cultures. Cells were induced at 0.8 O_D600_ with 1 mM IPTG and grown for 3.5 hours at 37° C. Cultures were spun down at 3000 g for 15 minutes and snap frozen. Frozen pellets were then resuspended in 50 mM NaCl, 50 mM HEPES-NaOH pH 8.5 to 1 M NaCl, 50 mM HEPES-NaOH pH 8.5 with 5 mM BME and protease inhibitor (Roche 11836170001). Resuspended pellet was then heated at 80°C for 20 minutes and sonicated for 1 minute 1 s on/1 s off at 60% power before centrifuging for 15 minutes at 30000 xg. Supernatant was retained, and 50% w/v saturated ammonium sulfate was added for 1 hour at room temperature while nutating. This was spun down again for 15 minutes at 30000 xg, and this time the pellet was retained. Pellet was resuspended in DI water with 4 mM DTT and .22µm filtered, then purified on Macro Prep High S (Bio-rad 12009271) column with a gradient of 50 mM NaCl, 50 mM HEPES-NaOH pH 8.5 to 1 M NaCl, 50 mM HEPES-NaOH pH 8.5. Resulting fractions with tau were collected and nutated with 4x volume acetone containing 5 mM DTT overnight to isolate protein monomer. Tau was spun down the following day and resuspended in 100 mM NaCl, 10 mM HEPES-NaOH pH 7.4 with 5 mM DTT, before aliquoting and storing at -80°C. All tau was thawed, diluted, and treated with nucleases before use.

### RNase and DNase treatment of tau constructs

All constructs were treated with RNase and DNase before use to remove any nucleic acids that persisted during purification. K18 stock was diluted to 100 µM final volume. Reaction included 5 µL RNase I_f_ (M0243S) and 5µL DNase 1 (M0303) per 100 µL with associated reaction buffers diluted in 100 mM NaCl, 10 mM HEPES-NaOH, pH 7.4. Nuclease reaction buffers were diluted 10X for seeding assay and 100X for the in vitro aggregation assay. Reaction was allowed to proceed at 37°C for 15 minutes, before adding 5 μM EDTA and heating at 75° C for 20 minutes to inactivate nucleases. Tau was placed on ice for 10 minutes before homogenizing and pipetting into assays.

### Seed formation

K18 or K19 was diluted to 25 μM in 100 mM NaCl, 10mM HEPES-NaOH pH 7.4. Respective polyanions were added to 12.5 μM on a per strand basis. Seeds were incubated on thermomixers at 37°C for 3 days at 550 rpm shaking. Seeds were removed after 72 hours and placed on ice for 15 minutes, then sonicated for 20 seconds in a water bath before use. Adapted from [13].

### K18 transfection of HEK293 cells

Tau RD P301S FRET Biosensor cells (CRL-3275 ATCC) were grown and maintained in DMEM with 10% Fetal Bovine Serum, 1% penicillin-streptavidin and 1% l-glutamine. Cells were plated at 300 μL per well in Ibidi (80826 8-well ibitreat) plates. 25 μM monomer K18 was incubated for 3 days, shaking at 37°C with 12.5 μM of the respective polyanion and RNase inhibitor (NEB M0314S) in 100mM NaCl, 10mM HEPES-NaOH, pH 7.4 buffer to create seeds. K18 was transfected into cells with 25 μL reaction mix and 25 μL lipofectamine mix. The reaction mix consisted of K18 and Optimem, while the Lipofectamine mix consisted of Lipofectamine 3000 and P3000. Each well received 20 μL of 25 μM K18, 1.5 μL lipofectamine 3000 and 1.5μL P3000 (Thermo L3000008). After 24 hours, cell media was removed and wells were incubated with 300 μL 1μ/mL Hoechst (CAT) in 1X PBS for 20 minutes. Hoechst was then removed and 300 μL fresh media placed into the wells. Images were taken at 1% intensity. ImageJ quantification was done using a puncta counting image analysis tool utilizing a Gaussian Blur background subtraction, minimum puncta size 0.5 μm^2^ to maximum puncta size 10 μm^2^, threshold method Otsu dark. https://doi.org/10.5281/zenodo.17662103.

### Thermal aggregation and spin down of 0N4R tau

25 μM K19 was incubated for 3 days, shaking at 37 °C in 100 mM NaCl, 10 mM HEPES, pH 7.4, with 12.5μM RNA and RNase inhibitor (NEB M0314S). Aggregation assays were performed by adding either 5, 10, or 20 μM RNA and 1 μM K19 seeds to 10 μM 0N4R monomer before shaking overnight at 37 °C. Adapted from [40].

### Electron microscopy

After an overnight aggregation assay of 0N4R with K19 seeds, samples were imaged by Transmission Electron Microscopy via negative staining. 10 μL of 5 μM 0N4R tau was pipetted on discharged copper grids for 5 minutes. Grids were then washed with water and incubated with 2% Uranyl acetate for 10 seconds before drying.

### Mass photometry

0N4R tau was diluted to 500 nM in triple-filtered 10mM potassium phosphate, pH 7.4 buffer. RNA was added at a 1:2 RNA:tau, 1:1 RNA:tau or 1.5:1 RNA:tau ratio. Samples were incubated for 30 minutes at room temperature before pipetting into grid wells on Refeyn CC Mass glass. Standards were either BSA (Sigma A7906), Beta-amylase from sweet potato (NATE-0762), or both. Molecular weights are displayed where they were given by the gaussian peaks fit through the Refeyn DiscoverMP peak-fitting software. Stoichiometries are estimated based on the following RNA molecular weights:

G4neg-RNA – 6.4 kDa, RNA-seq J19 – 7.4kDa, RNA-seq J6 – 7.4kDa, RNA-seq J23 – 6.6kDa, RNA-seq J20 – 8.2kDa

## 3. Results

### Nucleic acids enriched in human brain aggregates within the hippocampus form G-quadruplexes

We first sought to verify that potential G4-forming sequences found within hippocampal aggregates actually form G4s *in vitro*. We selected an initial set of the 22 enriched nucleic acid sequences with the highest likelihood of forming G4s based on bioinformatic predictions. [15] G4 formation was tested via circular dichroism (CD) (Fig 1A, Supplemental Fig. 1) and NMM binding (Supplemental Fig. 2). Four sequences displayed CD traces characteristic of parallel G4s, one formed an anti-parallel structure, and the rest were hybrid in structure with differing levels of parallal and anti-parallel topology. (Fig. 1A, Supplemental Fig. 1). Only one sequence did not appear to form a G4. (Supplemental Fig. 1). Many of the G4s formed oligomers with themselves when electrophoresed on a non-denaturing gel (Supplemental Fig. 1). Having determined that the vast majority of these sequences could form G4s as predicted, we set out to identify whether the G4s could modulate protein aggregation.

DNA and RNA G4s can act as effective chaperones for modulating protein aggregation and catalyzing protein folding. [23, 25, 26] We therefore tested whether these brain aggregate-enriched DNA G4 sequences possessed chaperone activity. We began by testing the effect of the 22 DNA sequences on the aggregation of citrate synthase upon its denaturation by heat. All G4s showed some effect on the protein aggregation when compared to the wells containing protein alone. Five G4 sequences almost entirely eliminated the formation of higher-order aggregates. (Fig 1C) Parallel and mixed G4s generally exhibited greater chaperone capability, while antiparallel G4s did not (Supplemental Fig. 2), consistent with previous observations. [25, 36] These data showed that many of the G4s enriched in human brain aggregates have powerful aggregation modulation properties, in this case, preventing aggregation of citrate synthase.

We chose the two sequences that almost completely eliminated citrate synthase aggregation under heat stress for further analysis. We used right angle light scattering to monitor oligomerization or aggregation using chemically denatured protein. We previously showed that in these assays, nucleic acids can kinetically stabilize protein-nucleic acid networks with nucleic acids separating partially folded protein oligomers, increasing the solubility of denatured proteins, and that many G4s do so very efficiently. [25, 38]

Citrate synthase protein was chemically denatured overnight and then diluted into a buffer that permits aggregation either alone or in the presence of the chosen G4s. Both sequences prevented the aggregation of denatured citrate synthase. The kinetics of the initial phase of light scattering with two DNA G4s were significantly faster than protein alone but plateaued with less total protein aggregation. (Fig. 1D) We have previously investigated this behavior and shown that it is indicative of G4 catalyzing the formation of small partially folded protein-nucleic acid oligomers held together by nucleic acid networks. [25, 38] These results suggested that the physiological G4s modulate protein aggregation by promoting small protein oligomer formation at the expense of greater higher-order protein aggregation.

### G-quadruplexes from brain aggregates modulate tau fibril formation

We then asked how these sequences might behave with a protein that is implicated in protein aggregation disease, such as tau. Tau monomer remains highly soluble under harsh conditions *in vitro*, but forms higher order fibril structures in multiple diseases *in vivo*. We set out to determine how these physiological G4s might affect fibril formation.

For *in vitro* experiments, we used the 0N4R tau construct, which contains a highly positively charged region previously shown to mediate RNA binding at physiological pH and the microtubule-binding domain of tau. We subjected this construct to heat stress and agitation with the addition of various RNAs and truncated seeds. We selected 3 DNA sequences from our initial experiments to test as RNAs. Two of these sequences had opposing activity in the citrate synthase thermal aggregation assay: RNA-seq J19 prevented all higher order aggregation of citrate synthase, while RNA seq J6 exhibited very little activity. We also added RNA seq J23, as it is a transcript from the APOE gene, and we previously found a significant correlation with APOE allele type and G4 formation in the human hippocampus. [35] We compared activity using an adenine-rich control RNA of similar length that does not form G4, hereafter termed G4neg-RNA.

Circular dichroism confirmed that all 3 RNA sequences formed primarily parallel G4s and that the control G4neg-RNA, as expected, did not fold into a G4 (Fig. 2A). Both G4 RNA sequences J19 and J23 showed monomer and higher-order oligomers as visualized on a native gel, while G4 RNA seq J6 did not. (Supplemental Fig. 5).

**Figure 2.**
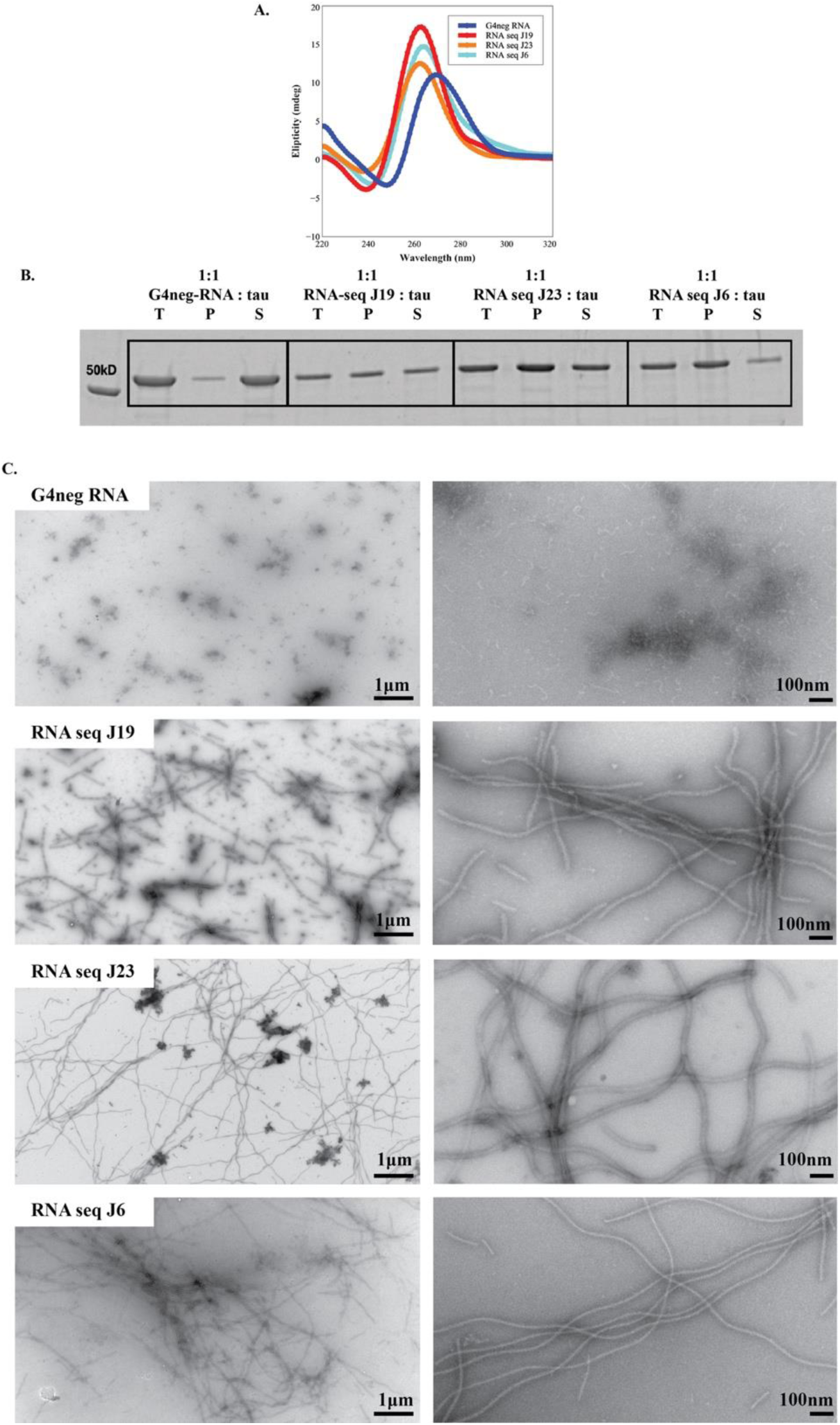
A) Circular dichroism of RNA sequences shows RNA seq J19, J23 and J6 form parallel G-quadruplexes, while the control RNA does not fold into G4. B) Spin down assay of 0N4R with G4neg-RNA, and G4 sequences indicates a higher population of insoluble tau with the addition of G4s. C) Electron microscopy confirms all 3 G4s form fibrils, but with to a varying extent and exhibiting morphological differences. No fibrils appeared with the addition of only the control RNA.

0N4R tau remained almost completely soluble with the addition of only seeds and the G4neg-RNA. (Fig. 2B) Upon the addition of each of the three G4 sequences at the same concentration, the solubility of tau decreased, and tau appeared in the pellet as well. (Fig. 2B) The G4 sequences varied in activity level, but all increased the amount of tau found in the insoluble fraction, especially RNA-seq J6. (Fig. 2B)

We then asked whether this insoluble fraction was forming tau fibrils. When examined via negative stain Transmission Electron Microscopy (TEM), the 0N4R tau formed fibrils with the addition of the G4s, but not with the G4neg-RNA. (Fig. 2C) All 3 G4s tested exhibited differences in tau fibril morphology and length. (Fig. 2C) RNA seq J19, which stabilized smaller oligomers with citrate synthase in a concentration dependent manner, promoted the formation of shorter tau fibrils. RNA seq J6, which did not stabilize smaller oligomers or prevent protein aggregation with citrate synthase, promoted the formation of longer, intertwined fibrils. More complicated higher order structures formed by many bunched up fibrils appeared in several wells with RNA-seq J6. These structures were in the tens of micrometers long. (Fig 2C, Supplemental Fig. 6) This was consistent with RNA-seq J6 exhibiting the greatest insoluble shift of tau in the spin-down assay. RNA seq-J23 also promoted longer fibrils, but these differed in appearance. These fibrils consisted of more individual strands with less bunching, wider diameter and more opaque center. The G4neg-RNA showed no fibril formation, consistent with the spin-down assay observations. This combination indicates the G4s interact with 0N4R tau in a different manner than the G4neg-RNA, allowing them to promote fibrillar states of tau, but also in ways that are distinct depending on the specific G4.

### RNA G-quadruplexes seed tau inclusions in cells

Since these brain-aggregate enriched G4s were very potent at forming tau fibrils *in vitro*, we hypothesized that they may be able to seed tau aggregation in cells despite being significantly shorter than those previously used in similar cellular tau assays. [14]

To test the effects of different RNAs in seeding tau inclusions, we utilized a cell line expressing K18 tau, the four microtubule binding repeats, with a disease mutation P301S. The cell line expresses a K18 fluorescent pair that forms fluorescent puncta when K18 is brought together in aggregates. We incubated recombinant monomer K18 tau with RNA-seq J19, RNA-seq J23, or G4neg-RNA for 3 days at a 1:2 RNA-to-K18 tau molar ratio at 37 °C while shaking. This pre-incubated K18 was then transfected into cell lines, and the number of fluorescent puncta was counted as a readout for seeding capability.

The G4neg-RNA promoted minimal formation of fluorescent puncta, comparable or less than protein without any polyanion. However, the G4 sequences RNA-seq J19 and RNA-seq J23 caused the formation of robust fluorescent puncta spread throughout the cells. (Fig 3A,B) This data shows that the G4s facilitate seeding and puncta formation in cells. As this assay started from monomeric tau, we therefore questioned whether G4s could themselves cause tau to form small oligomers.

**Figure 3.**
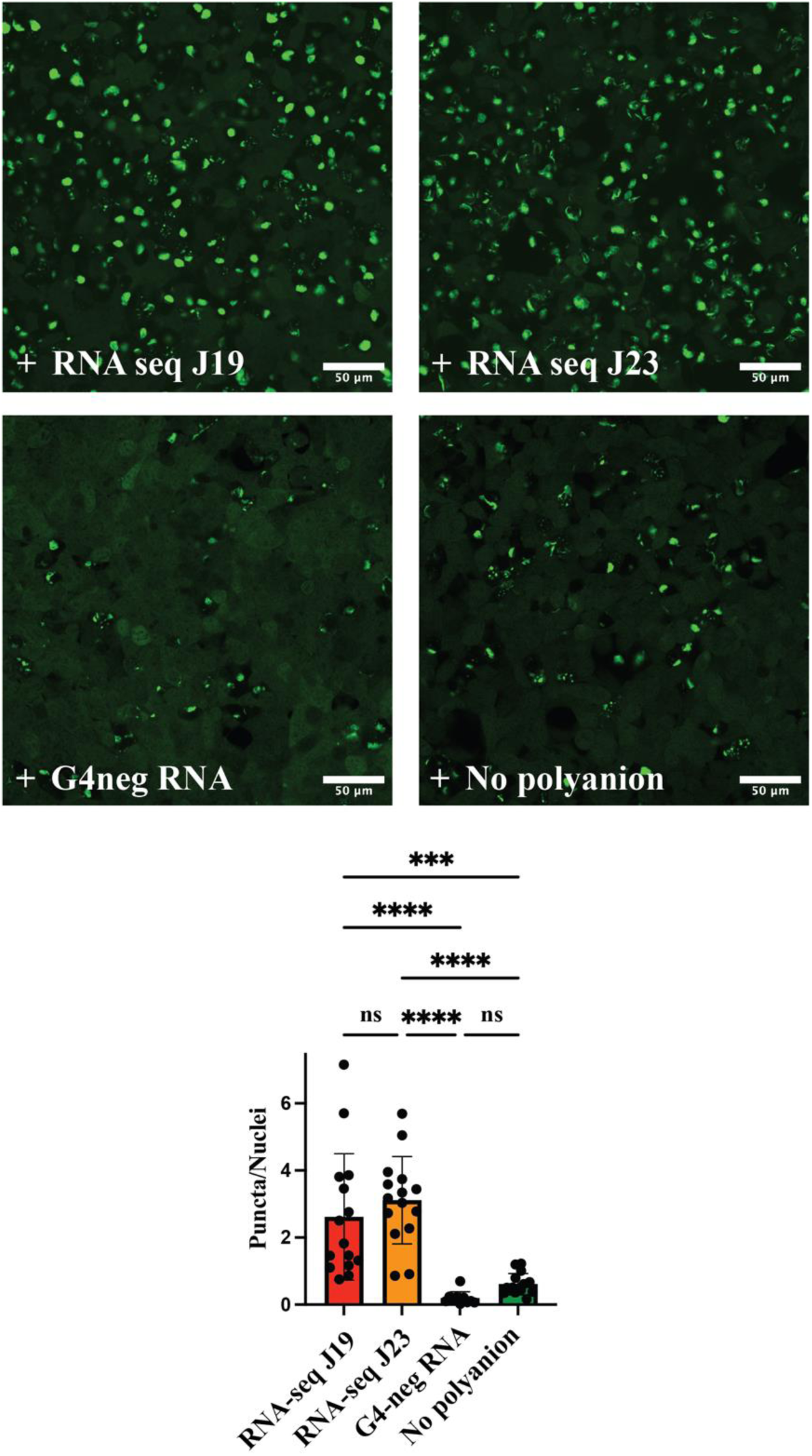
A) Fluorescent puncta appear with the addition of K18 incubated with G4neg-RNA, RNA seq J19 or RNA seq J23. K18 incubated with control RNA and no polyanion seed significantly fewer punctae. B) Image quantification of the number of fluorescent puncta. Significance represented by the ANOVA test with Tukey’s multiple comparison test. Error bars = SEM. Data shown represents quantification of three biological replicates, five images each.

### RNA G-quadruplexes promote different oligomer states

Since these G4s were particularly potent at generating seeding-competent tau, we questioned whether they may be sufficient to shift tau oligomer states on their own, without the addition of seeds. We tested the 0N4R construct via TEM with only the addition of the G4 RNA, revealing heterogenous populations of smaller tau oligomers. (Fig. 5A) We hypothesized that the G4s could bridge the binding of multiple tau monomers in a way that unstructured RNA does not. To test this hypothesis, we utilized mass photometry, a single-molecule technique that uses contrast to determine the mass of oligomers in solution.

0N4R tau alone was observed to form a primarily single peak in solution at around the predicted monomer size of 49 kDa (Fig. 4B), sometimes accompanied by a minimal dimer peak (Supplemental Fig. 8,10,11) Upon the addition of G4neg-RNA at a molar ratio of one RNA to two tau, we observed a shift of the monomer peak to higher molecular weight consistent with one RNA binding to one tau. In the presence of G4neg-RNA, no dimerization or further oligomerization of tau was observed, indicating that single stranded RNA was able to bind to but not oligomerize tau. (Fig 4C, supplemental Fig. 9,10,11)

**Figure 4.**
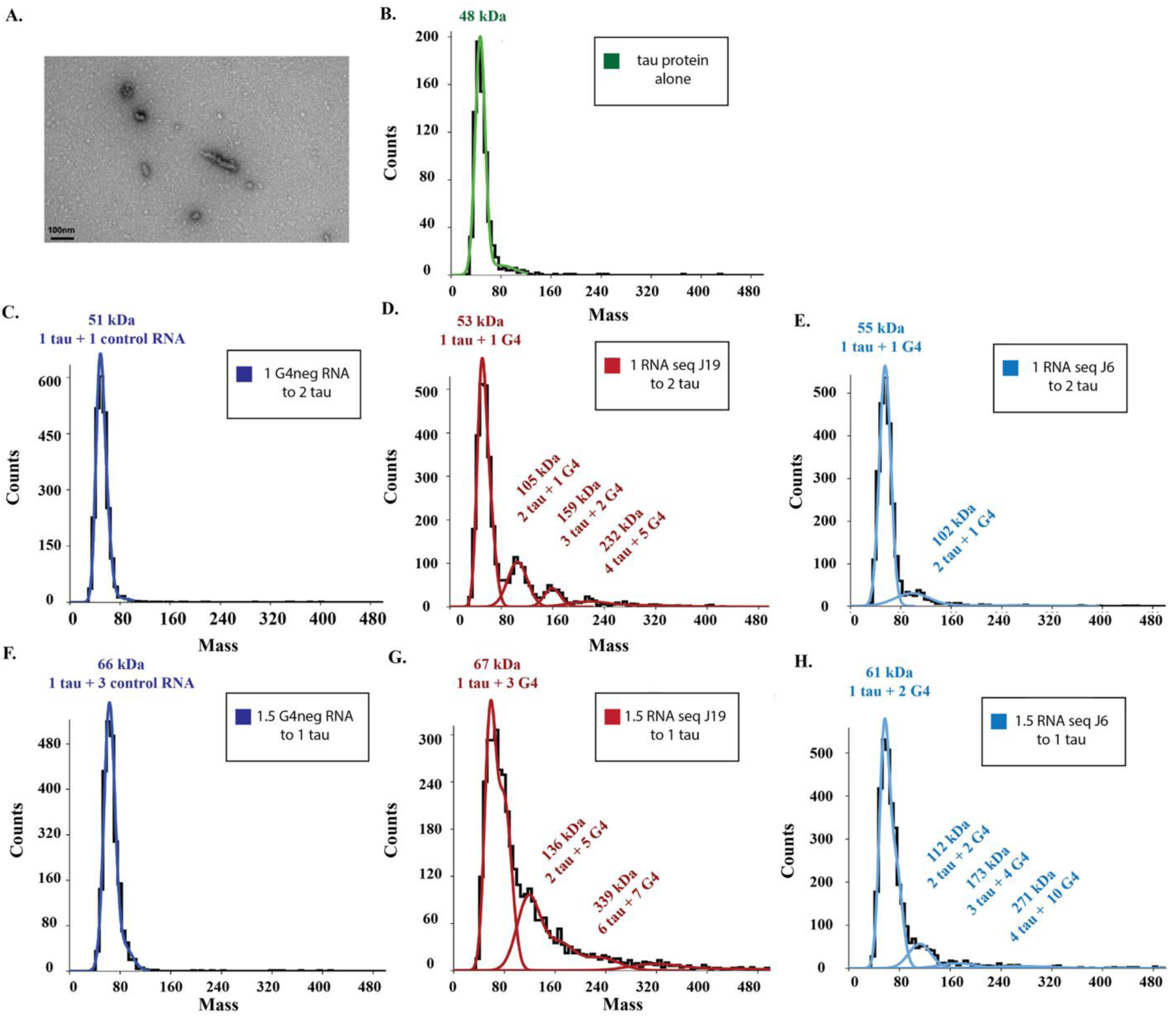
A) TEM images reveal various stabilized oligomer states with 0N4R tau and G4 RNA sequences alone, without the addition of seeds. B) 0N4R tau alone remains mostly monomer. C) G4negRNA does not change oligomer state of tau. D) G4 sequence RNA seq J19 exhibits a significant shift into distinct oligomer populations. E) G4 sequence RNA seq J6 shifts tau to mostly dimer population. F) Increasing concentration of G4neg-RNA increases binding of RNAs to tau monomer, but does not alter oligomeric state. G) RNA seq J19 shifts tau into higher oligomer states, with an increasing tail above 250kD. H) RNA seq J6 shifts tau into higher-order oligomeric states

We then examined three G4s with variable activity in the tau aggregation assay. For G4 RNA-seq J19, the effects on oligomer formation were striking. Upon the addition of this G4 sequence at a molar ratio of one tau to one G4, we observed binding to the monomer peak similar to our observations with the G4neg-RNA seq. Surprisingly, this sequence also caused the formation of three additional peaks, indicating the formation of higher-molecular-weight oligomers. (Fig 4B, Supplemental Fig. 10) These peaks were consistent with two tau bound to one G4, three tau and two G4s, and four tau and five G4 molecules, although other oligomer populations are possible as well for the highest molecular weight oligomer species. These results reveal that this G4 sequence can shift tau to distinct higher-order oligomer states at low concentration, without the need for seeds.

G4 RNA-seq J6 exhibited more RNA binding to the monomer and shifted tau into a dimer state consistently, but to a lesser degree than RNA-seq J19. Increasing the concentration of seq J6 relative to tau also revealed three additional independent populations, consistent with either two, three or four tau molecules bound to varying amounts of G4 molecules. (Figure 5E, 5H) G4 RNA-seq J20 either shifted the tau into two differently sized populations with a long tail towards higher molecular weights and less distinct peaks, or monomer and dimer with an even longer tail. (Supplemental Fig. 8) Both seq J6 and seq J20 exhibited more variability than the populations of seq J19. These results showed that at stoichiometric concentrations, G4s can cause tau oligomerization on their own, and that the oligomer states formed by tau depend on the specifics of the G4 sequence.

When we increased the concentration of the RNAs, we observed an increase in the molecular weight of the monomer peak for both G4 and G4neg-RNA, indicating multiple RNAs binding to the one tau. (Fig. 5G) Once again, the tau incubated with G4neg-RNA remained only one tau bound to multiple RNA molecules, while the G4s shifted tau into higher molecular weight species with multiple tau molecules bound. The G4 sequences caused a broader distribution of tau oligomer species at higher concentrations, with a significant increase in species above 200kDa. These higher concentration experiments show that the effects of the G4s are concentration dependent and highlight that the level of enrichment of these G4s could play a role in determining their overall effects *in vivo*. The unique ability of these G4s to shift tau into higher order oligomers appears to be a rare feature amongst other RNAs typically used with tau, suggesting they may play an outsized role in diseases involving protein aggregation and RNA dysfunction.

## Discussion

In this study, we examine the effects of nucleic acids enriched in human brain aggregates. These enriched sequences were previously predicted to form G4s, and we found that they largely did so *in vitro*. These enriched G4s exhibited strong effects on protein oligomerization and aggregation *in vitro* and in cells, at lengths and concentrations considerably shorter and lower than seen for other types of nucleic acids. However, the G4s had different effects depending on their sequence.

We observed that G4s generally accelerated oligomerization and prevented higher order aggregation of citrate synthase, and caused oligomerization and fibril formation of tau. For tau, different G4s caused changes in small tau oligomer populations, protein solubility, and contributed to intracellular aggregation to varying degrees. TEM and mass photometry revealed that G4s modulate tau oligomer populations, without the addition of tau seeds. All G4 sequences examined shifted oligomer states, but to drastically different oligomer sizes and heterogeneity. Our data shows this activity varies significantly even between three parallel quadruplexes, and does not correlate directly with guanine content, suggesting the sequence and topology specificity is more complex than previously thought.

Although previous work found that a G4 could shift tau into immobile phases more efficiently than other RNA sequences like PolyA, the RNAs examined were model sequences selected and likely not of physiological relevance. [11] The resulting aggregates appeared amorphous when observed under TEM, unlike the physiologically relevant G4 sequences we tested here that could promote fibril formation. This difference reflects the importance of continued studies to identify the specific physiologically relevant G4 sequences in different diseases, as they do not all have the same exact effects.

Much work has observed tau monomer conversion in the presence of polyanions such as heparin or polyA RNA, along with the addition of seed-competent tau. [13] However, questions remain on the physiological relevance of these *in vitro* studies on RNA, in part due to the high concentrations and large size of these RNAs. [14, 41–43] These studies typically utilize bulk poly(A) RNA or other long homopolymers, ranging from hundreds to thousands of nucleotides. Previous cell work reveals a minimum length dependency at around 40 nucleotides for seeding assays. [14] Shorter oligonucleotides induce LLPS, [11, 12] but longer fibril formation with 20 nucleotide RNA has not been shown. This highlights how powerful these short G4s studied are at oligomerizing and promoting tau fibril formation by comparison.

It was not only surprising that RNAs this short could induce longer fibril formation, but also that these fibrils varied considerably in structure. RNA-seq J19 stabilized shorter fibrils as opposed to RNA-seq J6. RNA-seq J19 fibrils were not smooth or continuous, while RNA-seq J6 induced fibril formation that were continuous for tens of micrometers long with symmetrical higher order structures. The morphological differences were striking considering all three sequences are parallel G4s, indicating their activity cannot be attributed to topology alone. RNA-seq J19 is composed of 77% guanines, RNA-seq J19 is 60% guanines, RNA-seq J23 is 55% guanines, RNA-seq J6 is 68% guanines. The guanine content did not correlate with seeding activity in the cells, extent of fibril formation, or mass photometer shifts. Therefore, protein interactions are governed not only by topology or G-richness, as previously assumed.

On a technical note, mass photometry has previously been utilized with the 0N4R tau construct also used here and showed monomer, dimer, and trimer peaks for tau alone, unlike in our case where nearly entirely monomer was observed. [44] The difference can be explained by our protocol including a monomer isolation step to ensure we are starting from the monomer state, in addition to a nuclease treatment prior to the assay, which removes residual nucleic acids remaining after purification. K18 tau without this nuclease treatment failed to induce puncta formation in HEK cells, despite a strong co-purified nucleic acid presence, indicating that not just any bound RNA will convert tau to a seed-competent form.

Previous studies showed that G4s fold with stress and apparently unfold when stress is removed. [21] We previously showed that G4s increase in hippocampal tissue with the chronic stress of age and disease severity, and high co-localization of G4s to phospho-tau aggregates. [35] Here, we observed that the concentration of G4s in relation to tau strongly affected fibrillation depending on the G4 sequence. Combined with the results presented here, these observations suggest that higher occurrence of certain G4s, or changes in the folding levels of these G4s, may play a role in tau aggregation in disease.

Overall, the RNA component of aggregates is more complicated than previously thought, exhibiting drastically different aggregate composition based on the structure and sequence. This observation emphasizes the importance of identifying which G4 sequences are involved in diseases and studying them separately in detail for their individual contributions. These data suggest a mechanism of G4-protein interaction and roles in oligomerization that is considerably more complicated than the typical charge-neutralization mechanism proposed previously for other nucleic acids. Moreover, the differences observed in oligomerization could impact other important and currently largely unanswered questions on protein aggregation, such as the variability between diseases with the same misfolded proteins but different fibril morphologies.

**Table 1.**
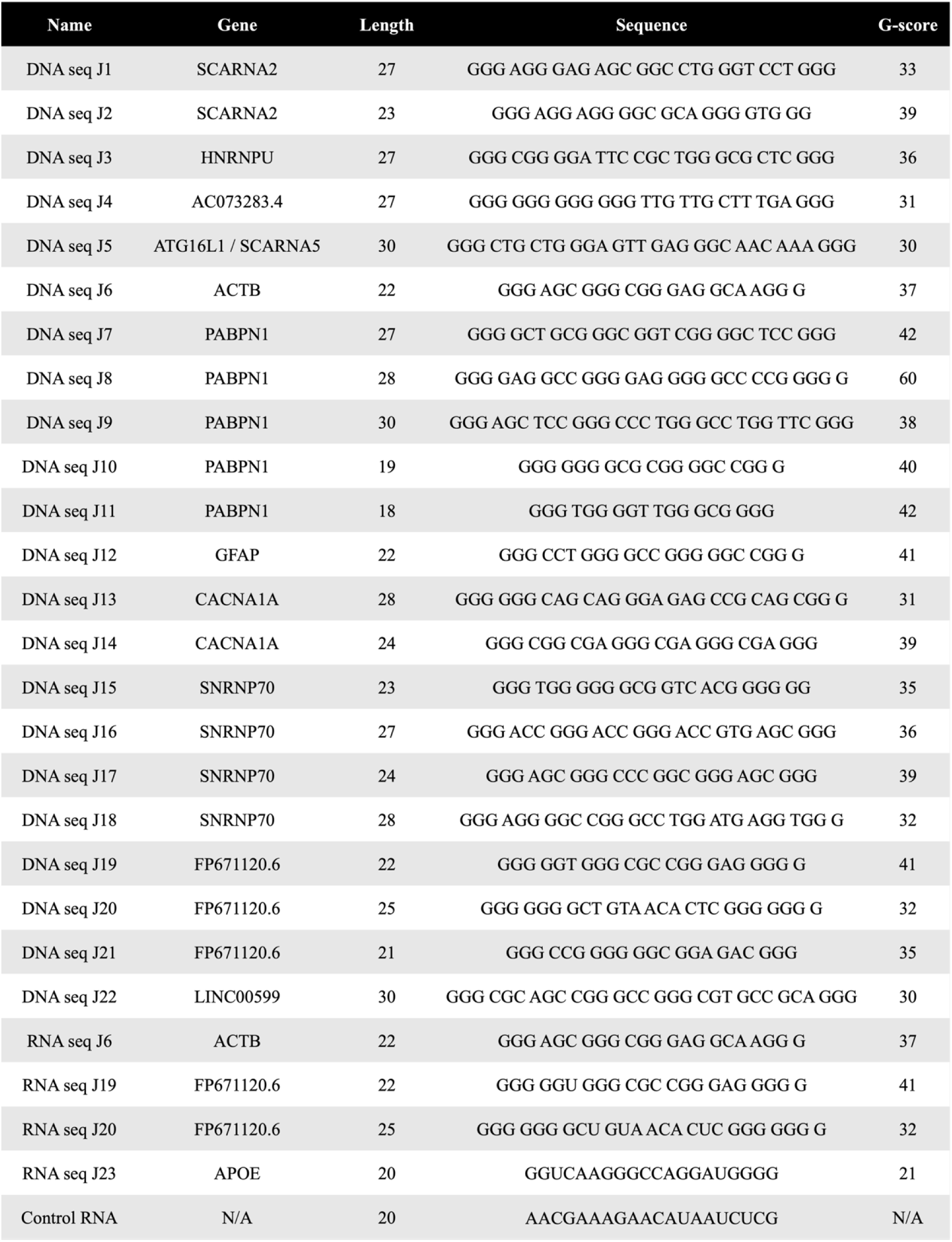
Nucleic acid sequences used in this study. From Table S4b G4 seqs. [15].

## Acknowledgements

Funding from NIH R35GM142442 to S.H. Images taken on JEOL JEM 120i TEM funded by NIH grant 1S10OD036258-01.We thank B. Sahoo and J. Bardwell for their helpful comments during editing. We thank M. Margittai for the K18 plasmid and test protein, and J. Bardwell for the K19 and 0N4R WT plasmid.

## Author contributions

Idea generated by S.H. Experiments designed by S.H. and L.M.K., with assistance from K.M.R and L.T.K. Experiments performed by L.M.K and L.T.K. Primary writing done by S.H and L.M.K., with editing and comments from all authors. Authors declare no competing interests.

